# Phenologs at knowledge-graph scale: a mouse-to-slime-mold catalog of candidate genes for thousands of human diseases

**DOI:** 10.64898/2026.10.23.761438

**Authors:** Aaron Odell, Daniel R. Korn, Bryan J. Laraway, Phillip A. Richmond, Nicolas Matentzoglu, Kevin Schaper, Christopher J. Mungall, Melissa A. Haendel

## Abstract

Model organisms carry decades of systematic gene-phenotype annotation, but connecting those annotations to human disease is rarely straightforward. Phenologs offer a principled way to bridge this gap. A human disease and a model-organism phenotype are phenologs when their associated genes, linked by orthology, overlap more than expected by chance, so the genes of one become candidates for the other. Here we compute phenologs from the Monarch Knowledge Graph, which integrates gene-disease, gene-phenotype and orthology data for human and seven model species. Using PANTHER protein families as orthogroups and a 5% false discovery rate estimated from 1,000 randomized networks, we identify 61,243 phenologs between 3,752 human diseases and 9,700 model-organism phenotypes. We introduce a hypergeometric scheme for ranking the candidate genes a disease’s phenologs nominate and compare it with the naive Bayes scheme of the original phenolog studies. Both methods recover held-out genes within a disease at similar rates, but only the hypergeometric score can be compared across diseases. Using a one-year-older release of the graph, the top 100 candidates per disease recovered 9 of 73 gene-disease associations added since, 23.5 times the random-gene expectation. Mouse accounts for most of these, with zebrafish, worm and fission yeast each contributing associations that mouse does not. We release per-disease gene candidate rankings, with known and novel associations flagged, and a dashboard for exploration of our results.

## Introduction

### Intro to gene-disease-phenotype

Gene-disease association remains a central challenge and unfinished task in biomedical research as thousands of Mendelian and complex diseases still lack a known molecular cause. Even when causal genes are known, the mechanisms linking genotype to phenotype are often opaque. Model organisms are a powerful tool for uncovering these mechanisms; decades of systematic mutagenesis and phenotyping in mouse, zebrafish, yeast, and many other diverse organisms, have produced rich gene-phenotype maps. However, transferring these model-specific insights to the realm of human disease is rarely straightforward. Conserved biological processes can manifest as seemingly unrelated phenotypes across species, while superficially similar phenotypes may arise from fundamentally different mechanisms.

### Intro to phenologs

The concept of phenologs, introduced by (McGary et al. 2010), offers a principled way to connect phenotypes across species. Two phenotypes from different organisms are termed “phenologs” when the genes associated with them, linked by orthology, overlap far more than expected by chance, signalling a shared underlying genetic mechanism even when the organism-level phenotypes may appear unrelated. Although the idea is framed initially in terms of phenotypes, the actual comparison taking place is more general. Gene-association sets derived from two different species can be compared, regardless of what the associations represent, as long as orthology between gene sets is known. In this study, each “phenolog” comparison pairs a human gene–disease set with a model-organism gene–phenotype set.

Because the relationship is defined by gene overlap rather than phenotypic resemblance, phenologs are mutually predictive: a gene known to participate in one phenotype of a pair becomes a candidate for the other. For example, previous phenolog analysis of *Arabidopsis thaliana* (mustard plants) nominated the gene SEC23IP as a candidate for the neural-crest defects of Waardenburg syndrome in humans (McGary et al. 2010), and the comparison of mouse and yeast gene-phenotype networks in the context of angiogenesis phenotypes motivated a drug repurposing screen that identified the antifungal thiabendazole as a vascular-disrupting agent, validated in frog and mouse models (Cha et al. 2012).

### What are our goals here?

In this work, we aim to improve upon the two previous studies from (McGary et al. 2010; Woods et al. 2013) along two axes to provide gene-disease candidate associations across a broad spectrum of human disease types. The first axis in which we aim to improve upon is the annotation substrate from which we derive our gene to phenotype network associations, and the second axis being the candidate gene-disease ranking mechanism post initial phenolog calculations. Prior works assembled gene–phenotype and gene-disease annotations from many individual resources, and computed cross-species gene orthology information per study using InParanoid (Ostlund et al. 2010). Here, we pull all annotations from the Monarch Knowledge Graph (Putman et al. 2023), a unified resource that harmonizes the necessary gene–phenotype, gene–disease, and orthology data under a common semantic data model (Unni et al. 2022). PantherDB protein families provide the orthology backbone (Thomas et al. 2022), and human disease terms are drawn from the Mondo ontology (Vasilevsky et al. 2025). In regards to gene-disease candidate scoring, pooling phenologs across a single disease can implicate many candidate genes but does not prioritize them, a problem that compounds as more species are combined. We introduce a hypergeometric (hg) scheme that weighs each candidate against its background frequency across the entire cross-species subgraph, and evaluate it against the prior method established in both previous studies.

### This work’s contribution summary

In all, we (i) compute phenologs across a wide breadth of human disease and phenotype terms using the Monarch KG at an estimated 5% false-discovery rate through an efficient, re-runnable procedure; (ii) introduce and evaluate a new scheme for gene-disease candidate ranking, showing it matches previous methods on recovery, improves precision, and enables candidate scores to be comparable across diseases; (iii) characterize how individual species contribute to human-disease predictions from the Monarch KG; and (iv) illustrate the framework through disease case studies, led by Bardet–Biedl syndrome. We release our results as per-disease candidate-gene rankings across 3,752 disease terms, with known and novel associations flagged together, as a resource for gene-disease prioritization. Together, this work updates the phenolog framework on a modern, knowledge-graph-scale dataset and demonstrates the Monarch KG as a substrate for cross-species disease-gene discovery.

## Methods

### Data gathering

We compute phenologs on data extracted from the Monarch Knowledge Graph (Putman et al. 2023), rather than attempting to query over the graph at scale. Leveraging the graph’s underlying Biolink data model (Unni et al. 2022), we are able to select for only the pertinent node and edge types across 8 different species including human, mouse, rat, zebrafish, frog, worm, fission yeast, and slime mold (**Fig. 1A**) using each species’ respective gene and phenotype ontologies. Gene orthology relationships are represented by edges derived from PantherDB (Thomas et al. 2022), with each edge carrying a protein family identifier, and inter-species gene-gene homology classification of ortholog, or least diverged ortholog (**Supplementary Note 2**).

**Figure 1.**
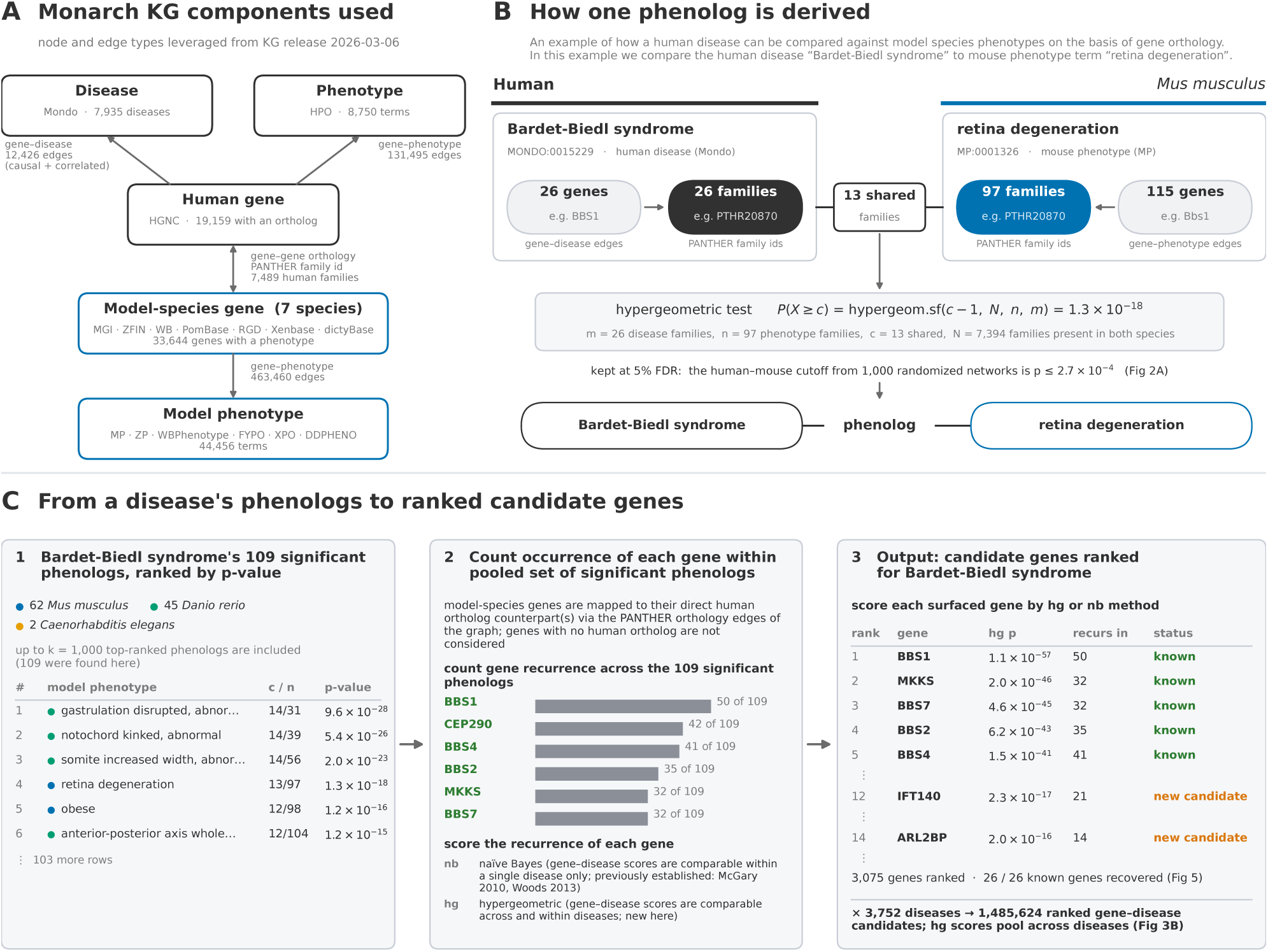
Phenologs formulation and results from Monarch KG. Overview of the phenologs methodology applied to the Monarch KG. (A) Components of the KG used in our analysis. Both human gene-disease, and gene-phenotype networks are linked by gene orthology (PANTHER) to 7 different species gene-phenotype networks. (B) Example of how a human disease term can be compared against a model species phenotype term through gene and protein family level orthology. PANTHER family ids are used to collapse genes to protein family level structure to scale calculations across uneven gene expansion/contraction between species. (C) How gene-disease candidates are scored across many associated phenotypes for a single disease (which can then be applied across all of the Mondo human disease ontology). Genes surfaced through guilt by association are tallied and pooled across all associated model phenotypes. Then each gene-disease candidate pair is ranked with respect to its peers within a single disease using the previously established naive Bayes method or using our new scoring method which allows for a score comparable both within a single disease and across all diseases. Note, we also perform the same pipeline for the human gene-phenotype network, but only the gene-disease example is shown for simplicity.

### Orthogroup assignment

Because gene families often expand and diverge unevenly across species (e.g. PTHR10574, which has only 1 gene in humans but 19 in mouse), computing the proper overlap of genes between any two species’ phenotypes is difficult to perform fairly at scale. Where (McGary et al. 2010; Woods et al. 2013) computed orthogroups with InParanoid we instead use the PantherDB protein family identifiers already integrated into each orthologous gene edge within the Monarch Knowledge Graph (**Fig. 1**). Collapsing genes to their PANTHER protein family gives a single cross-species unit for comparison. 19,159 human protein coding genes within the Monarch KG reduce to a set of 7,489 PANTHER protein families (**Figs. S1 and S2**), and 19,023 (99%) have a direct orthologous edge to a gene in at least one of the seven non-human species compared against.

### Phenolog calculations

Following data extraction, we take every pair of a human disease and a model-organism phenotype, and test whether their orthogroup sets overlap more than expected by chance using the upper-tail hypergeometric probability **(Fig. 1B)**. In the example given, we compare the disease Bardet-Biedl syndrome to mouse phenotype term retina degeneration by examining the overlap between each term’s respective known associated genes. The overlap is deemed significant if the p-value falls below a precomputed threshold determined by one thousand randomized trials, per set of species comparisons, aimed at controlling the false-discovery rate at 5% (McGary et al. 2010). Additional details can be found in (**Supplementary Note 3**). A significant overlap then nominates that phenotype’s genes as human candidates via guilt-by-association.

### False discovery rate filtration of initial phenolog calculations

The space of potential *(human disease, model organism phenotype)* pairs our algorithms can explore is large (#DISEASES * #PHENOTYPES) and if explored naively is prone to errors and excessive computation (**Supplementary Note 3, Table S4**). Therefore, we seek to perform a screening of unlikely results using the methods previously described by (McGary et al. 2010). The core idea of this filtration is that *“Biologically relevant disease/phenotype pairs in real data should have more overlapping orthogroups than would occur by chance.”* First, we create a distribution of scores for “non-biologically relevant relationships”, which takes the following three steps: ***(1)*** “*Shuffle*” relationships between human diseases to their orthogroups and model organism phenotypes to their orthogroups. ***(2)*** compute the significance scores for all pairs of these “randomized” disease-phenotype relationships. ***(3)*** repeat steps ***(1)*** and ***(2)*** 1,000 times and average the results to avoid biasing by any one specific randomized network. We then compare the “non-biologically relevant” distribution against the “true” distribution; finding the highest score such that there are 20x more “true” pairs than “fake” pairs (i.e. a 5% FDR) (**Fig. S3**). This is discussed more in depth in (**Supplementary Note 3**).

### Gene-disease ranking

Because a disease can inherit gene candidates from many phenologs across several species, a final step integrates this pooled evidence into one ranked list per disease (**Fig. 1C**). These candidates are nominated at the protein-family level, but because a single family can contain several human orthologs and or paralogs (42% of families do), we resolve them to individual genes using the graph’s gene-level ortholog edges, attributing each phenolog’s signal to the specific human ortholog(s) of the model gene that surfaces it.

We score the resulting gene-disease candidates with one of two schemes. The first is the naive Bayes (nb) scheme of the original phenolog work from (McGary et al. 2010; Woods et al. 2013), which scores a candidate by accumulating, across a disease’s top-k phenologs, the probability that it is implicated by each (**Supplementary Note 4, Eq. S1**). The second, introduced here, is a gene-level hypergeometric (hg) test that instead scores each candidate against how often it appears across the entire cross-species subgraph given the pooled set of genes found within the disease’s top-k associated phenologs (**Supplementary Note 5, Eq. S4**). The workflow outputs, for every human Mondo disease term and HPO term, a ranked candidate-gene list with known and novel associations flagged together. Note, that “known” and “novel” are with respect to the Monarch KG’s set of existing gene-disease edges (**Fig. 1B,C**).

### Evaluation and comparison metrics

Gene-disease candidate predictions produced by the phenologs pipeline are evaluated under two different paradigms. The first method involves comparing the gene-disease candidate predictions produced from an older version of the Monarch knowledge graph (one year prior) to newly introduced gene-disease pairs only found in a more recent version (i.e. can the method predict new/real biology). This method is discussed and explored further in (**Results, “Gene-disease candidate evaluation” and Supplementary Note 6**). The second method is a leave-one-out cross validation scheme adapted from (Woods et al. 2013) whereby known human gene-disease candidates are withheld, and evaluated on the basis of recoverability using the established pipeline. Additional details regarding this method are found in (**Supplementary Note 7**).

## Results

### Initial phenologs results

In **Fig. 2A** we plot the distribution of p-values for the comparison of human disease terms against mouse phenotype terms, along with the p-value distribution for 1,000 trials of the same comparisons, but with randomized networks according to (McGary et al. 2010). As expected, an enrichment in lower (i.e. more significant) p-values can be seen in the real data, and this same real-versus-random gap holds for every species examined (**Fig. S4**). The same methodology is applied to the human gene-phenotype network against mouse gene-phenotype network in **Fig. 2B**, and shows the same pattern across all model species (**Fig. S5**). Taking only the disease phenologs below each species’ FDR p-value threshold, yields 61,243 significant phenologs across 3,752 Mondo human disease terms and 9,700 model-organism phenotypes (**Fig. 2C**).

**Figure 2.**
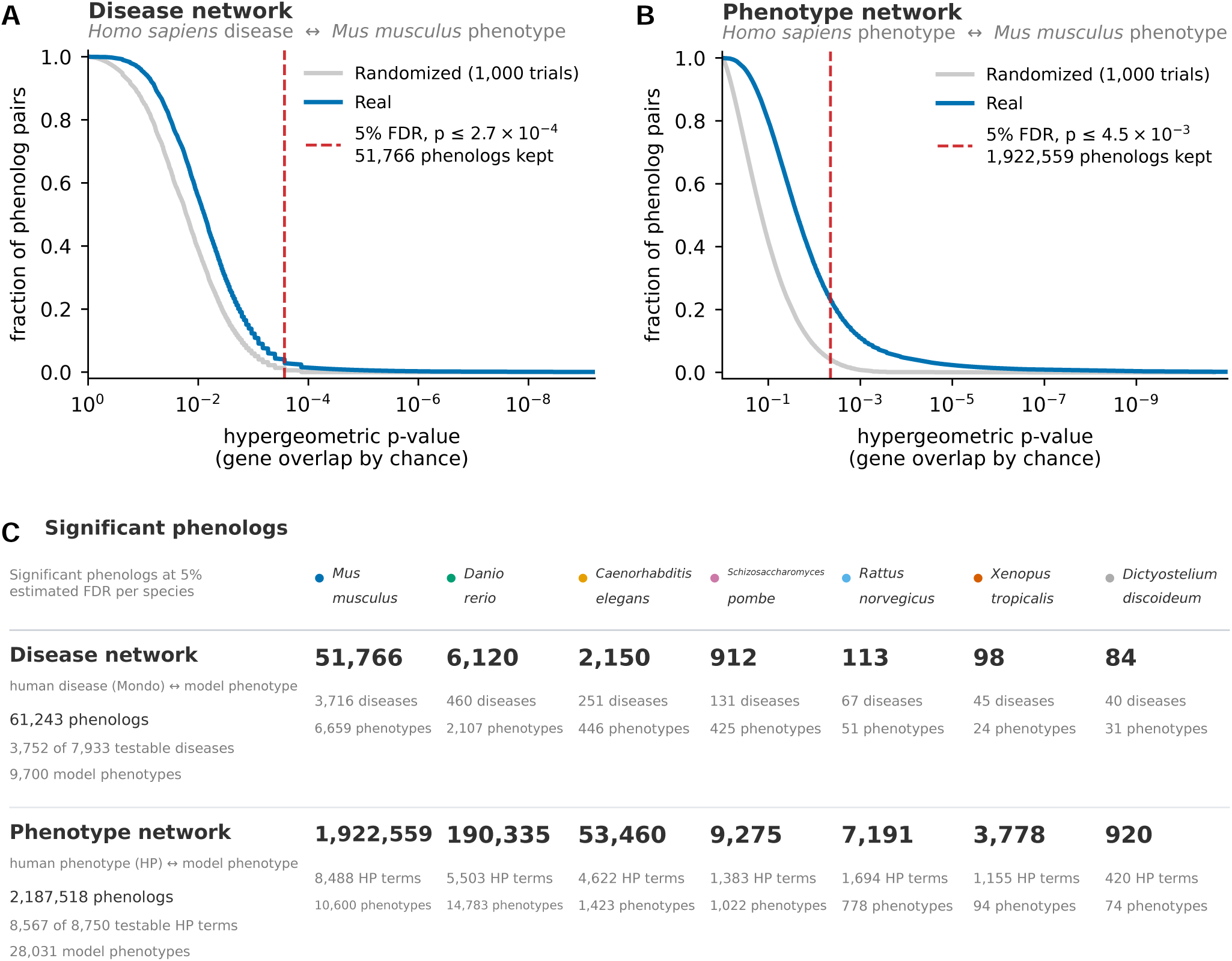
Phenolog detection at 5% estimated false discovery rate across two human networks and seven model species. (A) Results of the 1,000 trials between randomized human gene-disease and mouse gene-phenotype networks. Phenolog p-values are pooled across all 1,000 random trials and then sorted (for comparisons where there is ≥ 1 gene/ortholog overlapping). We highlight with the red dashed line the p-value representing an estimated 5% false discovery rate in accordance with work prior (McGary et al. 2010) (Methods, “False discovery rate filtration of initial phenolog calculations”). (B) Same as panel A, but for the comparison between human gene-phenotype network, and mouse gene-phenotype network. (C) Significant phenologs at 5% estimated false discovery rate for each model species, using the same methods described for panels A and B across all species, not just mouse. Each column represents a model species breakdown of the phenolog results obtained for the comparison against human gene-disease and gene-phenotype networks. The aggregates of all species results are tallied in the left most column. 61,243 phenologs surfaced over 3,752 diseases and 9,700 model phenotypes in the disease network comparison. The phenotype network comparison surfaced 2,187,518 phenologs over 8,567 HP terms and 28,031 model phenotypes.

This is more than an order of magnitude more human disease terms than earlier phenolog studies, a testament to more than a decade’s worth of progress since the original publication in 2010. Additionally, 3,088/3,752 (82.3%) of disease terms surfaced are the most-specific (leaf node) classes in Mondo, indicating genuine breadth of coverage across the disease ontology.

Examining the disease-phenologs surfaced across each model species shows the majority belong to mouse at 51,766 (84.5% of the total), but also shows meaningful signal is also drawn from the other species, including zebrafish (6,120 at 10.0%), fission yeast (912, at 1.5%), and down to just 84 for slime mold, tracking with each species’ initial annotation depth (**Table S4**). **Table 1** highlights the top two ranking disease phenologs found for each species we analyzed, with many associations appearing biologically cogent. For example, human retinitis pigmentosa pairs with a mouse phenotype for thin retinal outer nuclear layer, one of the most significant hits in our results, matching the known histopathology of the disease (O’Neal et al. 2024). While this example is intuitive in the semantic sense, an eye phenotype relates to an eye disease, the ability to obtain less semantically intuitive results is a core strength of the phenolog method.

**Table 1.** Top ranked disease phenologs for each species. Top two disease-phenotype associations from each species used in this study ranked by p-value significance. **m** = disease protein families, **n** = model phenotype protein families, **c** = shared protein families, **N** = protein families shared between human and model species, **p** = P(X≥c). Results demonstrate both semantically intuitive, and non-obvious phenolog results.

| Disease term | Cross-species phenotype | Species | m | n | c | N | hg p-value |
| --- | --- | --- | --- | --- | --- | --- | --- |
| hearing loss, autosomal recessive | increased or absent threshold for auditory brainstem response | *Mus musculus* | 61 | 211 | 42 | 7394 | 3.6e-52 |
| retinitis pigmentosa | thin retina outer nuclear layer | *Mus musculus* | 79 | 139 | 35 | 7394 | 7e-41 |
| Fanconi anemia | female sex determination decreased occurrence, abnormal | *Danio rerio* | 20 | 22 | 17 | 6479 | 1.7e-43 |
| Fanconi anemia | male sex determination increased occurrence, abnormal | *Danio rerio* | 20 | 24 | 17 | 6479 | 2.3e-42 |
| colorectal cancer | vulval cell induction increased | *Caenorhabditis elegans* | 21 | 12 | 6 | 3931 | 9.6e-12 |
| familial thoracic aortic aneurysm and aortic dissection | small | *Caenorhabditis elegans* | 12 | 48 | 7 | 3931 | 1.9e-11 |
| AP-4 deficiency syndrome | decreased protein localization to late endosome | *Schizosaccharomyces pombe* | 4 | 5 | 4 | 2301 | 4.3e-12 |
| hereditary breast ovarian cancer syndrome | abolished break-induced loss of heterozygosity via chromosomal translocation | *Schizosaccharomyces pombe* | 9 | 8 | 5 | 2301 | 1.3e-11 |
| chromosome 1p36 deletion syndrome | decreased fetal weight | *Rattus norvegicus* | 13 | 3 | 3 | 7296 | 4.4e-09 |
| postsynaptic congenital myasthenic syndrome | impaired spatial learning | *Rattus norvegicus* | 9 | 6 | 3 | 7296 | 2.6e-08 |
| cleft lip/palate | abnormal tail morphology | *Xenopus tropicalis* | 12 | 41 | 5 | 6173 | 7.7e-09 |
| Loeys-Dietz syndrome | abnormally ventrally curved tail | *Xenopus tropicalis* | 4 | 12 | 3 | 6173 | 2.2e-08 |
| undetermined early-onset epileptic encephalopathy | delayed aggregation | *Dictyostelium discoideum* | 27 | 30 | 6 | 3021 | 1.4e-07 |
| nevus, epidermal | aberrant pinosome maturation | *Dictyostelium discoideum* | 2 | 2 | 2 | 3021 | 2.2e-07 |

Indeed, **Table 1** demonstrates a link between the disease Fanconi anemia to skewed male-to-female offspring ratio in zebrafish, interpretable only through the shared DNA-repair pathway, in which loss of the gene FANCL drives Tp53-mediated germ-cell apoptosis and sex reversal (Rodríguez-Marí et al. 2010). We also created a dashboard to help visualize and explore results for both disease-phenologs and phenotype-phenologs https://monarch-initiative.github.io/monarch-phenologs/.

### Gene-disease candidate evaluation

To test whether our methodology can anticipate gene-disease associations that are absent from the graph entirely, we used the versioned history of the Monarch Knowledge Graph as a natural time split. We rebuilt the phenolog and gene ranking catalogs using a graph version released one year prior to our current graph version. We then assess how many of the newly added gene-disease associations within the current version of the graph (2026-03-06) are predicted by the phenolog calculations from the previous graph version.

Within the 1-year interval between graph versions, the gene-disease associations change only modestly (**Table 2**). About 96% of human gene–disease pairs are shared between the earlier and current release, with 524 total new pairs added. These added pairs are the raw material for our evaluation, but they are not uniformly usable, and scoring all of them together would skew the results. We therefore grouped them into categories to select for a reasonable evaluable set. Most new gene-disease pairs (402/524) fall on diseases that had no associated genes in the earlier graph, and are therefore untestable by our method, because a phenolog ranking needs a prior signal for the disease to build on (a known limitation when using in this way). Of the testable remainder, a substantial portion is redundant within the ontology. In other words, the exact pair is new, but the same gene was already linked in the earlier graph version to a more general, more specific, or sibling Mondo term for a similar disease. For example the gene MLH1, already tied to Lynch syndrome, reappearing under the sub-type of Lynch syndrome 1. In the context of evaluating the predicted gene-disease pairs put forward here, these redundant pairs likely reflect Mondo refinement rather than new biology, and are therefore not included in our analysis, resulting in a conservative 73 new gene-disease pairs available for evaluation.

**Table 2.** Gene-disease pair differences between Monarch KG versions. Breakdown of new gene-disease pairs added between Monarch KG versions separated by 1 year. New gene-disease pairs are categorized in order to assess relevance in evaluating phenolog gene-disease predictions. 73 new pairs were found where the candidate pair is non-redundant within the ontology between versions, and also the disease in the pair has at least 1 existing gene association already.

| Human gene-disease pairs (vs 2026-03-06) | 1 year prior (2025-03-10) |
| --- | --- |
| Shared by both versions | 11,902 |
| Newly added | 524 |
| Removed since prior | 335 |
| Genuine new link (evaluation set) | 73 |
| Redundant within the ontology | 49 |
| Not testable (no prior genes on disease) | 402 |

We assess the candidate predictions in two complementary perspectives. First, we examine on a per-disease basis in terms of recall and fold change over choosing genes at random (**Fig. 3A**). Fold change over a random gene here is defined as the top-N recall divided by the recall expected when, for each disease, N genes are drawn at random from the U human genes carrying an ortholog edge to a modeled species, equal to min(N, U)/U per gene–disease pair. It measures the enrichment of the full procedure, both restricting each disease to a phenolog-derived candidate set and ranking that set, over blind selection from the eligible gene universe.

**Figure 3.**
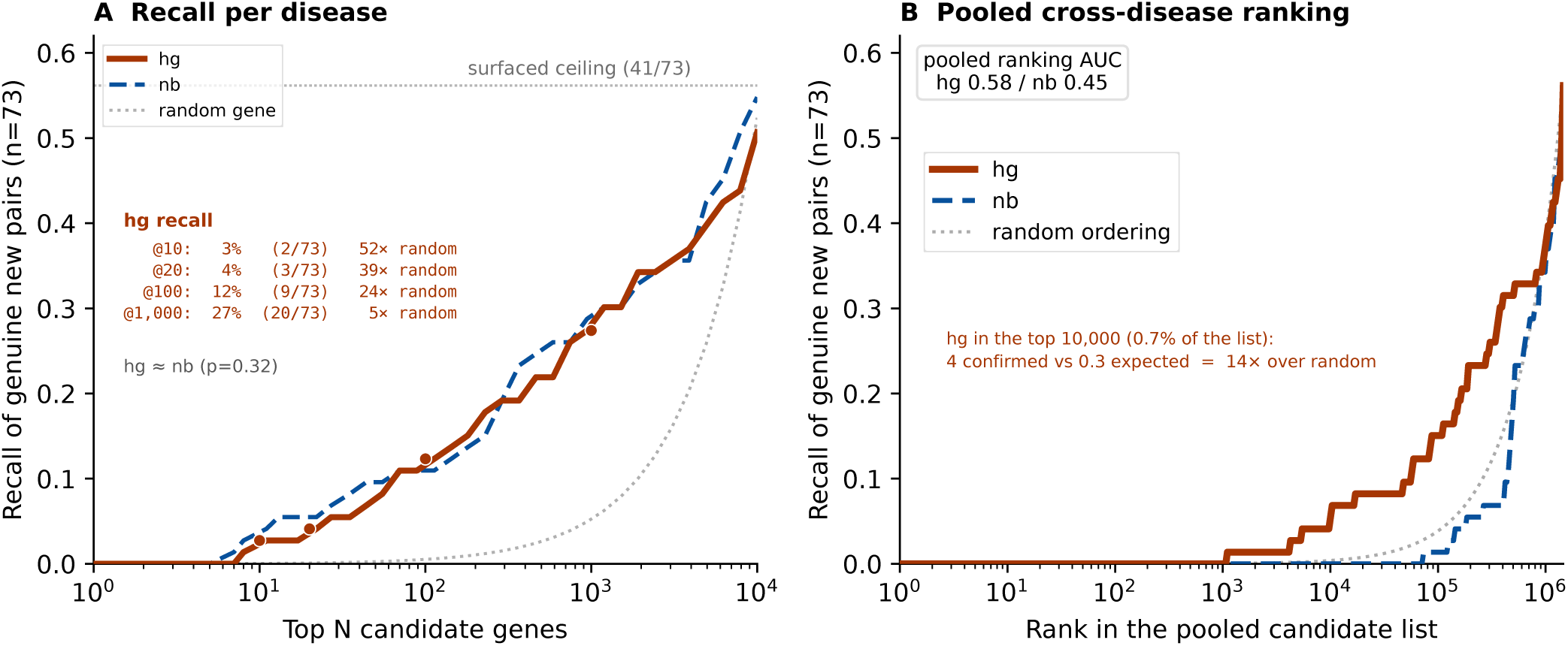
Temporal validation across graph versions. Gene rankings built from a 1 year prior graph version 2025-03-10, evaluated against the genuine new gene–disease pairs added by 2026-03-06 (n = 73; Table 2). All rankings use k = 1000 phenologs per disease. (A) Per-disease read: recall of genuine new pairs within each disease’s top N candidates (solid, hg; dashed, nb; dotted, genes drawn at random from the orthology-eligible universe). Insets tabulate hg recall at fixed top-ranked candidate depths with fold enrichment over the random-gene expectation. The horizontal dotted line marks the surfaced ceiling, the fraction of genuine pairs present anywhere in the phenolog-derived candidate lists (41/73). hg and nb are statistically indistinguishable at one year (p = 0.32). (B) Pooled read: all candidate pairs from all diseases merged into one globally ranked list per scoring scheme; curves show cumulative recall of genuine pairs by global rank (dotted, the same candidate set in random order). Only the cross-disease-calibrated hg score separates from random ordering (pooled AUC hg 0.58 vs nb 0.45). Within the top 10,000 globally ranked candidates (top 0.7% of candidates), 4 were confirmed over 0.3 expected at random chance (14x).

Evaluating against the genuine new gene-disease pairs, the method performs well above chance at prioritizing pairs it has never seen. Ranking with the hypergeometric score recovers 9/73 (12.3%) of the newly added genes within each disease’s top 100 candidates. This far exceeds the chance of a random draw at a 23.5-fold enrichment (95% CI 10.5 to 39.2) with a permutation p-value of 2×10⁻⁴ (Phipson and Smyth 2010). For robustness, we also perform a parameter sweep across the number of top ranked phenologs included per disease (k), and the top N candidate genes considered (**Supplementary Note 6, Fig. S6**).

Ultimately, our work here is computing all pairwise comparisons of human disease terms with model species phenotype terms, resulting in 1,485,624 novel and known gene-disease pairs, across 3,752 disease terms. Therefore, for the second assessment method (**Fig. 3B**), we pool all gene-disease candidate pairs together and assess recall expected over random when drawing the top N ranking pairs across our entire predicted set. When all candidate pairs are merged into one globally ranked list, the hg scoring method concentrates the confirmed pairs well above chance (pooled AUC 0.58) while nb is close to random ordering (0.45; **Fig. 3B**).

Within the top 10,000 pooled pairs, 0.7% of the list, hg recovers four confirmed associations while being 14-fold over the random expectation, demonstrating the utility of the hg method as an alternative scoring metric to the previously established nb method. We do note that the nb scoring method was not originally developed with this kind of analysis in mind and is largely a foundational piece of work to the methods developed and applied here.

Further analysis of the recovered gene-disease pairs revealed only one recovered association that was genuinely prospective in nature. FAAP100 was absent from the 2025 graph, and the method ranked it among the top candidates for Fanconi anemia on the strength of mouse phenolog evidence alone, and the human FAAP100–Fanconi anemia link was first reported in April 2025, after our reference snapshot (OMIM #621258). FAAP100 is a subunit of the Fanconi anemia core complex, most of whose other members already carried human disease associations, so its recovery follows the same conserved-module logic the phenologs method relies upon.

We performed an additional set of evaluation experiments by adapting the orthogroup-based leave-one-out cross validation of (Woods et al. 2013), hiding each of the 2,001 annotated protein families in turn and asking whether its genes are recovered by the method (**Supplementary Note 7, Fig. S7**). When ranking candidate genes within a disease, hg and nb recover hidden genes at near identical rates, as expected since both draw on the same candidate lists (**Fig. S7A**). When the same rankings are read across diseases, asking where a hidden gene’s true disease falls among all diseases it is ranked for, only hg retains meaningful resolution, placing the true disease within the top 20 for 165 diseases against 12 for nb at k = 20 (**Fig. S7B**). On the complete model, hg also recovers a larger share of each disease’s known genes at every top candidate list size, and is more precise in 67% of diseases (**Fig. S7C,D**).

### Individual species contributions

In addition to improving how evidence is integrated across phenologs, we sought to quantify each species’ individual contribution to disease information, a question made more pertinent by the heavily lopsided gene–phenotype annotation counts seen across each species (**Table S4**). Using the same framework from the previous section, we investigate the individual, and combined effect of each species’ contributions in predicting the newly added gene–disease associations. We rebuilt the hg rankings from each model species’ phenologs in isolation on the one-year-prior graph (k = 1000 top phenologs included in gene scoring, as before) and scored each against the same 73 genuine new pairs (**Fig. 4A**). The rankings built from all seven species align closely with the mouse-only ranking within a single hit at every candidate list depth, while also reaching three additional pairs mouse alone was unable to recover. These results indicate that pooling data across all species can buy additional coverage while maintaining the same level of precision.

**Figure 4.**
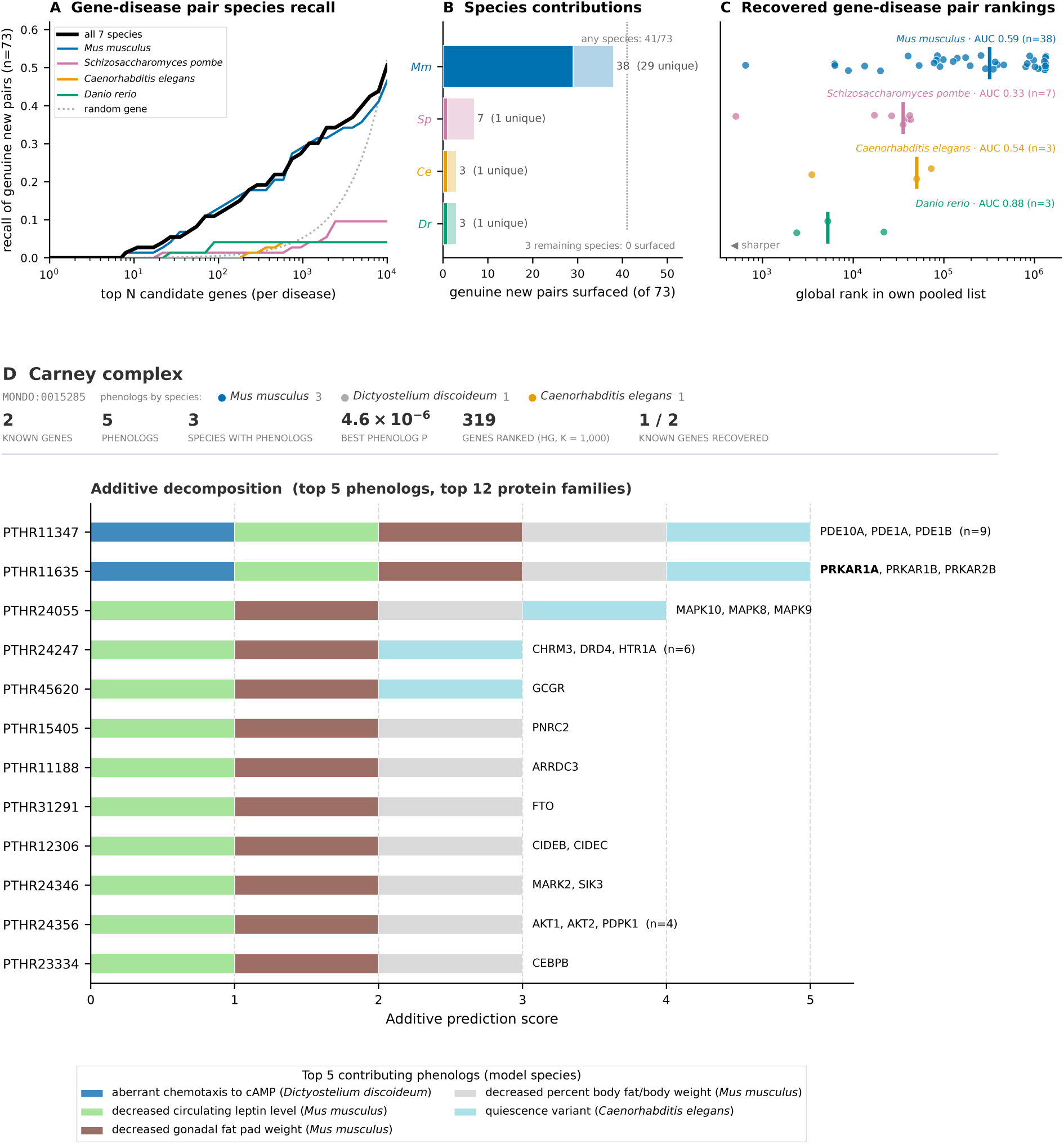
Species contributions to prospective prediction. Panels A-C use the evaluation results from Fig. 3: hg rankings built from the one-year-prior graph (2025-03-10; k = 1000 phenologs per disease), scored against the 73 genuine gene–disease pairs added by 2026-03-06. Species are abbreviated Mm, Mus musculus; Sp, Schizosaccharomyces pombe; Ce, Caenorhabditis elegans; Dr, Danio rerio. (A) Recall of genuine new pairs within each disease’s top N candidates, for the ranking built from all seven species combined and from each contributing species alone; dotted line, random genes. (B) Genuine new pairs surfaced in each species’ standalone candidate list (dotted line, the 41 of 73 reached by at least one species); solid segments are pairs unique to that species. The three remaining species (Rattus norvegicus, Xenopus tropicalis, Dictyostelium discoideum) surface none: their standalone lists contain at most 1,331 candidate pairs, versus 43,598 or more for the contributors, so the test is structurally blind to them. (C) Global rank of each reached pair within the species’ own pooled cross-disease list (log scale; vertical bar, median; AUC computed over that list). The fission yeast value is dominated by one broad disease term accounting for six of its seven hits; the seventh ATM ranks in the top 1.2% at a rank of 509 of 43,598. (D) Additive decomposition of Carney complex candidate gene rankings (MONDO:0015285). The header lists the significant phenologs by species (three Mus musculus, one Dictyostelium discoideum, one Caenorhabditis elegans) and summary counts: 2 known genes, 5 phenologs, 3 species with phenologs, the best phenolog p-value (4.6 × 10⁻⁶), 319 genes ranked by hg at k = 1,000, and 1 of the 2 known genes recovered in that ranking. Bars show the twelve top-scoring PANTHER protein family ids with representative gene symbols and the syndrome’s known surfaced gene in bold.

**Fig. 4B** shows the individual per species breakdown of the recovered gene-disease pairs. Of the 73 genuine new pairs, 41 are reachable by at least one species ranking alone. Mouse reaches 38 of them, 29 uniquely; fission yeast reaches seven, and worm and zebrafish three each, and each of these species contributes at least one pair that no other species reaches. In **Fig. 4C**, we further quantify the rankings of the recoverable 73 gene-disease pairs with respect to each species’ surfaced candidates. Mouse is broad but diffuse, carrying most of the coverage at moderate per-hit precision (AUC 0.59), while zebrafish is sharp but narrow, placing its three hits within the top 6.5% (AUC 0.88). Fission yeast appears worse than random (AUC 0.33), but six of its seven hits belong to a single broad disease term, complex neurodevelopmental disorder, reached only through generic growth and viability phenotype terms, and ranked accordingly low (between the 39th and 99th percentiles). The remaining recovered gene-disease pair from fission yeast links the gene ATM to hereditary breast ovarian cancer syndrome, and lands in its top 1.2% of predicted genes, aligning with fission yeast’s textbook use as a model organism for the study of DNA-repair (Humphrey 2000; Hoffman et al. 2015). The remaining three species (rat, frog, and slime mold) surface none of the newly added 73 gene-disease pairs, but this appears to be a structural blind spot of the analysis itself, rather than a judgement on their lack of signal. Both the low number of newly added gene-disease pairs from the latest graph version, along with low annotation counts for these species, leads to a low probability of intersection. However, this does not discount the value these species may add in modeling human disease in this way.

To further explore each species’ ability to model an aspect of human disease biology, we investigated the top ranking phenologs for each species using the results computed from the most recent version of the knowledge graph. Using the most evolutionarily distant species from human in our dataset, Dictyostelium discoideum, we provide an example in the context of the disease Carney complex, an endocrine tumor syndrome caused by the gene PRKAR1A (**Fig. 4D**). We use the additive gene candidate scoring scheme of (Woods et al. 2013) to decompose and visualize the candidate gene rankings for Carney complex by their associated phenologs. Each gene score is the sum of per-phenolog contributions and therefore can decompose into a stacked bar showing which phenologs, and species support each candidate on a linear scale, revealing each species’ contributions in modeling the cAMP–PKA module. Five significant phenologs were found in total: three mouse adiposity phenotypes (decreased circulating leptin, gonadal fat pad weight, and percent body fat), the Dictyostelium discoideum aberrant chemotaxis phenotype, and a Caenorhabditis elegans quiescence phenotype. The three mouse phenologs do support the two causal protein families, the phosphodiesterases and the PKA regulatory subunits containing PRKAR1A, but they support eight other families just as strongly, mostly fat and metabolism related families such as FTO and CEBPB. On mouse evidence alone, the causal module is buried in generic adiposity biology. The two invertebrate phenologs help resolve this tie. The quiescence phenolog adds support to a handful of signaling families, while the chemotaxis phenolog supports only the phosphodiesterase (PTHR11347) and PKA regulatory subunit families (PTHR11635), leaving them as the only two families backed by all five phenologs and placing them alone at the top of the ranking in our method. This is an example of how leveraging multiple species can fill in the gaps across the knowledge base, as well as a demonstration of the phenologs method itself.

Taken together, the contributions from each model organism largely overlap, but also are able to cover holes in the knowledge base no one single species is able to cover, with mouse showing the broadest coverage overall. We provide an additional example in the next section in the form of an in depth analysis of Bardet-Biedl syndrome where the phenologs method performs well, multiple species contribute to the signal, and where candidate-gene prioritization is still a clinical need.

### Phenolog analysis of Bardet-Biedl syndrome

Bardet-Biedl syndrome (BBS; MONDO:0015229) is an autosomal-recessive ciliopathy affecting roughly 1 in 160,000 people, and up to 1 in 13,500 in some genetically isolated populations. It is characterized by retinal degeneration, obesity, post-axial polydactyly, renal anomalies, hypogonadism, and cognitive impairment (Forsythe and Beales 2013). At least 24 genes encoding the BBSome, chaperonin, and intraflagellar-transport machinery are now implicated (Niederlova et al. 2019), yet even comprehensive sequencing of the known BBS genes resolves only ∼80% of clinically diagnosed cases, leaving roughly 20% without an identifiable pathogenic variant, a gap that has persisted across panel and exome cohorts as the gene set has grown (Forsythe and Beales 2013; Perea-Romero et al. 2022). Several further layers of complexity complicate the genetic picture: potential pathogenic variants in genes not yet linked to BBS, copy-number variants that elude targeted panels (Lindstrand et al. 2016), additional rare alleles in other ciliary genes that may modulate expressivity (Perea-Romero et al. 2022), and, in a residual fraction of patients, causes that remain unknown. Prioritizing candidate genes, both undiscovered and modifiers, is therefore not purely academic for BBS; it is part of the bottleneck between a clinical diagnosis and a molecular one.

BBS is well suited to cross-species inference because ciliary biology is deeply conserved and the syndrome is modeled particularly well outside of mammals (Masek et al. 2022). Of its 109 significant phenologs, 45 derive from zebrafish, whose photoreceptor and ciliary phenotypes closely mirror the human disorder, alongside 62 from mouse and 2 from worm. In **Fig. 5** we display the 20 top-scoring orthogroups over BBS’s 20 strongest phenologs, recovering most of the known BBS genes, jointly driven by mouse and zebrafish. Notably, the top phenologs such as retinal and photoreceptor degeneration, obesity, absent sperm flagella, and abnormal pronephric-tubule morphology are all recognizably BBS-like and recapitulate the ocular, metabolic, reproductive and renal features of the human syndrome (Forsythe and Beales 2013). By this same logic, genes surfaced alongside the known BBS panel are credible candidates for follow up investigation. We note, 13/20 top families are shared across all three gene scoring integration methods (nb, hg, additive), with hg and additive both recovering 15/20 top families, and nb at 16/20. In all, the phenologs methodology and analysis presented here is seemingly able to model Bardet-Biedl syndrome particularly well.

**Figure 5.**
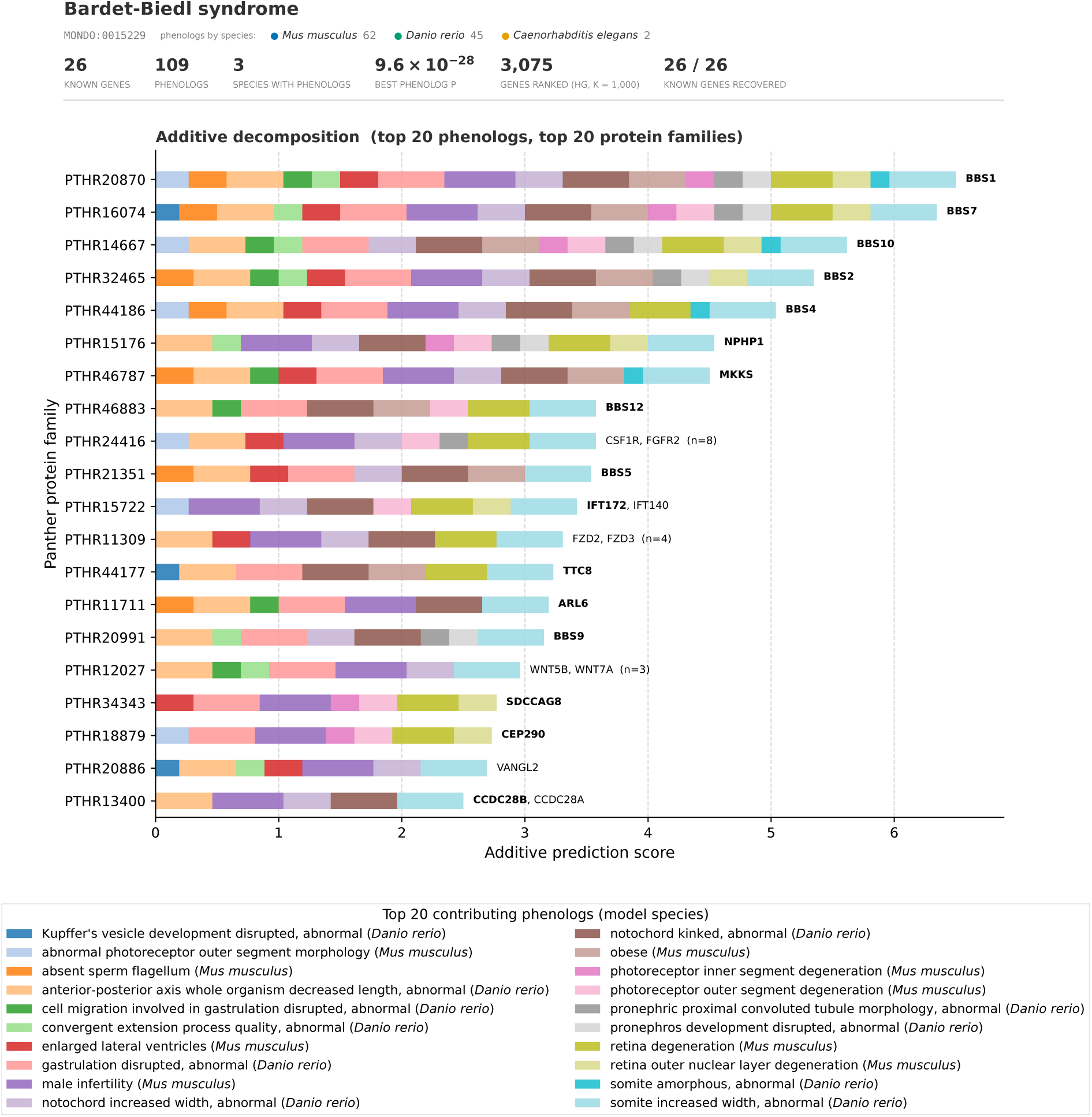
26/26 known gene-disease genes for Bardet-Biedl syndrome are recovered. Top 20 protein families surfaced by the top 20 most significant phenologs for Bardet-Biedl syndrome. The header gives the disease identifier (MONDO:0015229), the significant phenologs by species (62 Mus musculus, 45 Danio rerio, 2 Caenorhabditis elegans) and summary counts: 26 known genes, 109 phenologs, 3 species with phenologs, the best phenolog p-value (9.6 × 10⁻²⁸), 3,075 genes ranked by hg at k = 1,000, and 26 of 26 known genes recovered in that ranking. Every horizontal bar is one PANTHER protein family, and its length is that family’s prediction score decomposed additively into the contribution of each supporting phenolog. The human genes in each family are listed at the right, with the family’s total gene membership count in parentheses. Genes already associated with Bardet–Biedl syndrome are shown in bold. Each contributing phenotype term and its model species is shown in the legend. 16/20 top-scoring protein families correspond to established Bardet–Biedl genes (bold). BBS1, BBS2, BBS4, BBS5, BBS7, BBS9, BBS10, BBS12, MKKS, ARL6, TTC8, CEP290, SDCCAG8, IFT172 and NPHP1. The phenologs driving these genes span the cardinal organ systems of this disorder: ocular, metabolic, reproductive, and renal. Note, CCDC28B is also bolded because it is annotated to Bardet-Biedl syndrome subtype 1 (MONDO:0008854)

## Discussion

### Summary of our contributions

In this study we show that the Monarch Knowledge Graph, mined through the phenolog framework, yields candidate gene–disease predictions across a wide breadth of human disease, and we provide an extensive evaluation of how those predictions can be scored. Building on the candidate-gene prioritization task introduced by (McGary et al. 2010; Woods et al. 2013), we introduce a hypergeometric (hg) scheme for integrating gene-disease as an alternative to the original naive Bayes method (nb). We demonstrate the benefit of the hg method when ranking gene-disease evidence across the entirety of the predictions that are made (1,485,624 candidate pairs across 3,752 disease terms and 15,170 genes), an assessment the previous nb method was unable to perform. Additionally, under the leave-one-out methodology previously established, hg matches nb when ranking candidates within a disease, is the only scheme of the two that remains informative when candidates are compared across diseases, and improves precision at the top of each disease’s candidate gene list (**Fig. S7**).

### Value of cross-species data integration

We examine the benefits of cross-species genetic and phenotype data integration for human gene-disease prediction through worked examples of the phenologs methodology (Carney complex, and Bardet-Biedl syndrome), and a per species analysis of a temporally split set of gene-disease edges from the Monarch KG, demonstrating the ability of 4/7 species to predict edges not currently in the graph. Mouse, by virtue of its extensive gene-phenotype annotations compared to the rest of the species, supplies candidate genes across most disease areas. Pooling data from all model organisms surfaces an additional 3 gene-disease pairs unreachable when using only mouse annotations, while maintaining the same quality in ranking.

### Human disease biology modeled through the lens of evolutionary biology

An exemplar of the phenologs paradigm in its purest form is the analysis of Dictyostelium discoideum, a slime mold without a nervous system, an endocrine system, or a skeleton. This organism is extensively removed from human biology, with its divergence from the nearest human ancestor occurring approximately a billion years ago (Fiz-Palacios et al. 2013). Yet, studying this organism still provides insights into puzzles of our own biology.

### Phenologs catalog as a resource

The datasets produced here are intended as both a current resource, for human gene-disease and gene-phenotype association exploration and a building block for future work. We release the full set of disease-phenologs, and the top-100 candidate genes per human phenotype term (HP) as a set of parquet files https://monarch-initiative.github.io/monarch-phenologs/?network=disease#/downloads. The candidate-gene lists for the phenotype network are limited to the top 100 per HP term for file size convenience only. The most immediate application of this dataset is gene and variant prioritization for unsolved patient cases. Complex diseases such as Bardet-Biedl syndrome, where ∼20% of cases lack an identifiable variant in a known primary or oligogenic modifier gene (Shoemaker 2024), is where a ranked list of candidate genes and modifiers to interrogate could prove valuable. While we provided an extensive analysis of the human disease-phenolog dataset, we barely scratched the surface of the human phenotype-phenolog dataset. Integrating this information with patient phenotype profile data and optional genetic workup could offer value as orthogonal approach for unsolved cases. We also find it worth noting that in the age of extremely powerful AI systems both commercially available and open-source, novel hypothesis generation from datasets like these is trivial, even at scale. Evaluation, the ability to carry out experiments, and produce anything meaningful from such predictions is all much larger in scope, and is thus excluded from our analysis here.

### Limitations

The phenologs method requires known genetic associations to exist beforehand in the underlying networks being analyzed. This “seed” requirement before new gene associations can be inferred limits the human disease and phenotype terms that are reachable through this method alone. Human genes with no ortholog available in another species are also unreachable by this method as it is bound by evolutionary conservation. Lastly, not all available information has or is able to be ingested into the Monarch KG in regards to human gene-disease information and is an ongoing effort.

## Conclusion

Together, this work revitalizes the phenolog framework on a modern, knowledge-graph-scale dataset, and associates seven sets of model species gene-phenotype information to thousands of human disease and phenotype terms. More broadly, it demonstrates the Monarch Knowledge Graph as a substrate for discovery, turning decades of model-organism phenotyping, from mouse to slime mold, into prioritized hypotheses for the genetic basis of human disease.

## Data availability

The phenolog catalogs and candidate-gene rankings described here are distributed as release 2026-03-06 of the monarch-phenologs resource, built from Monarch KG release 2026-03-06 (PHENIO 2025-11-14). The disease-network release holds the 61,243 significant disease phenologs, the 1,485,624 ranked gene-disease candidates across 3,752 Mondo disease terms (hg score, k = 1,000, 5% FDR) with known and novel associations flagged, and the supporting gene, phenotype and orthology tables. The phenotype-network release holds the 2,187,518 phenologs across 8,567 HP terms with the top 100 candidate genes per term. Results are made available for both download and exploration at https://monarch-initiative.github.io/monarch-phenologs/. The Monarch KG releases used in this study (2025-03-10 and 2026-03-06) are available at https://data.monarchinitiative.org/monarch-kg-dev/.

## Code availability

The phenologs pipeline, dashboard, and the release tooling are open source under the BSD 3-Clause licence at https://github.com/monarch-initiative/monarch-phenologs. The results reported here were produced with commit 9cd1477 (2026-10-14); the public release files were exported with commit 294b8d7.

## Author contributions

Conceptualization: A.O., B.J.L.

Methodology: A.O., B.J.L., N.M., K.S.

Software: A.O., B.J.L.

Formal analysis: A.O., B.J.L.

Investigation: A.O., D.R.K., B.J.L., P.A.R.

Resources: A.O., N.M., K.S.

Data curation: A.O., B.J.L., N.M., K.S.

Writing – Original Draft: A.O., D.R.K.

Writing – Review & Editing: A.O., D.R.K., P.A.R., C.J.M., M.A.H.

Visualization: A.O., D.R.K., B.J.L., P.A.R.

Supervision: K.S., C.J.M., M.A.H.

Project administration: A.O., M.A.H.

Funding acquisition: C.J.M., M.A.H.

## Funding

This work was supported by the Office of the Director, National Institutes of Health under award numbers 1R24OD042574 (An AI-powered informatics resource for accelerating NAM research), 1R24OD039947 (An explainable AI platform integrating diverse data for enhancing rigor, reproducibility, and translatability of animal research) and 5R24OD011883 (The Monarch Initiative: linking diseases to model organism resources). The content is solely the responsibility of the authors and does not necessarily represent the official views of the National Institutes of Health.

## Competing interests

The authors declare no competing interests.

## Acknowledgements

We thank the Monarch Initiative team and the model organism database communities, including MGI, ZFIN, WormBase, PomBase, RGD, Xenbase and dictyBase, Mondo, and HPO, whose curated gene-phenotype, disease, and phenotype annotations this work depends on, along with the PANTHER, Mondo, and HPO teams for the orthology, human disease, and human phenotype resources.

## Supplementary Information

### Supplementary Note 1. Monarch KG filtering

The Biolink Schema is used by the Monarch KG as an underlying data model representation. We identified the relevant sets of nodes and edges for the phenologs calculations we wanted to perform and created a subgraph of the larger Monarch KG to begin our analysis.

**Table S1.** Monarch KG Biolink Categories, Associations, and Predicates. **Monarch KG node and edge types selected for.** The Biolink node and edge types we select for are listed within the table for our analysis. Note that GeneToGeneHomology edges were restricted to protein coding gene pairs where a PANTHER protein family classification is given as evidence within the edge provenance.

| Kind | Biolink category | Predicate |
| --- | --- | --- |
| Node | biolink:Gene | n/a |
| Node | biolink:PhenotypicFeature | n/a |
| Node | biolink:Disease | n/a |
| Edge | biolink:GeneToPhenotypicFeatureAssociation | biolink:has_phenotype |
| Edge | biolink:CausalGeneToDiseaseAssociation | biolink:causes;<br>biolink:associated_with_increased_likelihood_of |
| Edge | biolink:CorrelatedGeneToDiseaseAssociation | biolink:gene_associated_with_condition;<br>biolink:contributes_to |
| Edge | biolink:GeneToGeneHomologyAssociation | biolink:orthologous_to |

Additionally, each species within the knowledge graph uses its own ontologies for phenotype and or disease node name representation (**Table S2**). Depending on the structure of the ontology used, there may or may not be terms needed to be excluded from our analysis (we are primarily interested in only abnormal phenotypes). To achieve our filtering goals of only including gene-phenotype associations where an alteration of the “gene” is causing or likely causing an abnormal phenotype, we precompute sets of nodes from each ontology to either include or exclude from our initial subgraph building process. We reconstruct each species’ individual ontology from phenio, a larger overarching ontology linking many ontologies together that the Monarch KG leverages under the hood in each release (Putman et al. 2023). This allows us to achieve version parity with all ontologies used within a single extra file, rather than downloading each ontology individually (https://github.com/monarch-initiative/phenio/releases/latest/download/phenio-relation-graph.gz). Each ontology’s node/edge structure can then be reconstructed by following the directed graph from the edges linking between intra ontology nodes based on the prefix identifiers (e.g. HP, MP, MONDO). Filtering can then be performed based on whether or not a term descends from a chosen root term (**Table S3**). For example, fission yeast in our calculations only includes phenotype terms that are descendants of the “abnormal phenotype” term within the Fission Yeast Phenotype Ontology (FYPO) (Harris et al. 2013).

**Table S2.** Species disease and phenotype ontologies. **Ontologies used for each species within the Monarch KG.** We select specifically for nodes with these prefixes within the subgraph building process, along with the relevant biolink model category. For example, a node with an ID beginning with “ZP:” would also need to coincide with the biolink category PhenotypicFeature.

| Taxon label | Phenotype ontology | Disease ontology | Phenotype prefix | Disease prefix |
| --- | --- | --- | --- | --- |
| *Danio rerio* | Zebrafish Phenotype Ontology | n/a | ZP | n/a |
| *Xenopus tropicalis* | Xenopus Phenotype Ontology | n/a | XPO | n/a |
| *Caenorhabditis elegans* | C. elegans Phenotype Ontology | n/a | WBPhenotype | n/a |
| *Rattus norvegicus* | The Mammalian Phenotype Ontology | n/a | MP | n/a |
| *Mus musculus* | The Mammalian Phenotype Ontology | n/a | MP | n/a |
| *Schizosaccharomyces pombe* | Fission Yeast Phenotype Ontology | n/a | FYPO | n/a |
| *Dictyostelium discoideum* | Dicty Phenotype Ontology | n/a | DDPHENO | n/a |
| *Homo sapiens* | Human Phenotype Ontology | Mondo | HP | MONDO |

**Table S3.** Ontology term filtering. **Phenotype and disease term filtering via ontological structure.** Not all phenotypes are necessarily valid for this study and thus needed to be avoided when creating the initial subgraph of the original KG. We select for abnormal phenotypes, and human disease specifically when it is needed. Each ontology was examined visually using EMBL-EBI Ontology Lookup Service (McLaughlin et al. 2025) in order to identify the proper terms.

| Ontology | Terms included |
| --- | --- |
| Mondo | Only "human disease" terms (Include everything under MONDO:0700096) |
| Human Phenotype Ontology | Only "phenotypic abnormality" terms (Include everything under HP:0000118) |
| Fission Yeast Phenotype Ontology | Only "abnormal phenotype" terms (Include everything under FYPO:0001985) |
| The Mammalian Phenotype Ontology | Everything except for "normal phenotype" terms (Exclude everything under MP:0002873) |
| Dicty Phenotype Ontology | Everything except for "wild type" terms (Exclude everything under DDPHENO:0000142) |

### Supplementary Note 2. Gene homology

The primary source of gene homology information within the Monarch KG originates from PantherDB’s (Thomas et al. 2022) “AllOrthologs.tar.gz”, which catalogs all orthologous relationships between the gene sets of more than 140 different species. This information is ingested by the Monarch KG for a select set of species as gene-gene orthology relationships. Each gene with an ortholog in at least one other species is assigned a protein family identifier by PantherDB (e.g. PTHR11482). The eight species used in this study are the only species in which gene-phenotype relationships exist within the graph.

As described in (Woods et al. 2013), the direction of comparison between species-specific gene-phenotype networks becomes a computational enigma when homology is treated gene-by-gene, rather than collapsing orthologous and paralogous relationships into “orthogroups” that create a single point of comparison across divergent species gene families. We approximate this using the PantherDB protein family assignment for each gene, collapsing the 19,159 human genes carrying an assignment into 7,489 orthogroups and removing the asymmetry caused by unevenly expanded gene families.

**Figure S1.**
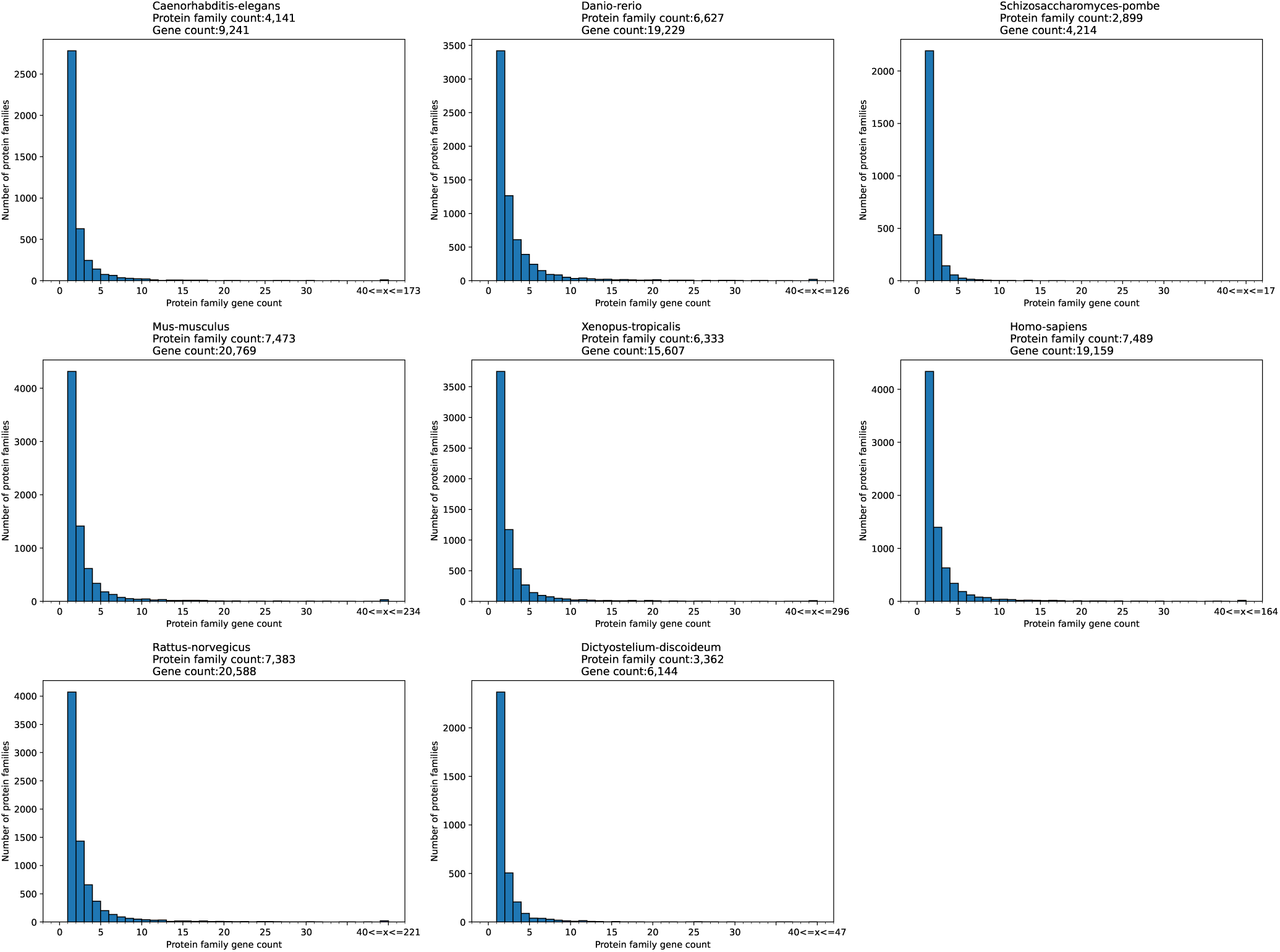
Distribution of the number of genes per protein family for each species used. PantherDB protein family gene count distribution for each species. We note this is a construct of the species representation within the Monarch KG in the sense that if a gene does not have an ortholog in another species within our dataset, it is not included. For example, by PantherDB’s count there are 19,450 protein coding genes in humans, but our dataset only captures 19,159 (98.5%) of them because of the species diversity limitation within our data and genes unique to humans.

We collapse at PantherDB’s family level (e.g. PTHR11482) rather than its finer subfamily subdivisions (e.g. PTHR11482:SF7), because the family level maximizes the shared cross-species members available to each comparison; subfamilies, or requiring a direct one-to-one ortholog edge, would fragment these sets and reduce the statistical power of the initial phenolog test. For this reason the orthogroup abstraction is used only to detect phenologs. However, the underlying gene-level ortholog edges are retained to rank gene candidates downstream. Each family-level signal is resolved back to the specific human ortholog(s) of the driving model gene (42% of human families contain more than one human gene).

The effects of this collapse on human disease nodes, in terms of gene and protein-family annotation counts, are shown in **Fig. S2**. The gene networks remain robust when collapsed to the orthogroup level: of the 1,116 human diseases annotated with two or more family-bearing genes, only 72 (6.5%) collapse to a single orthogroup, so multi-gene diseases almost always retain more than one point of comparison.

**Figure S2.**
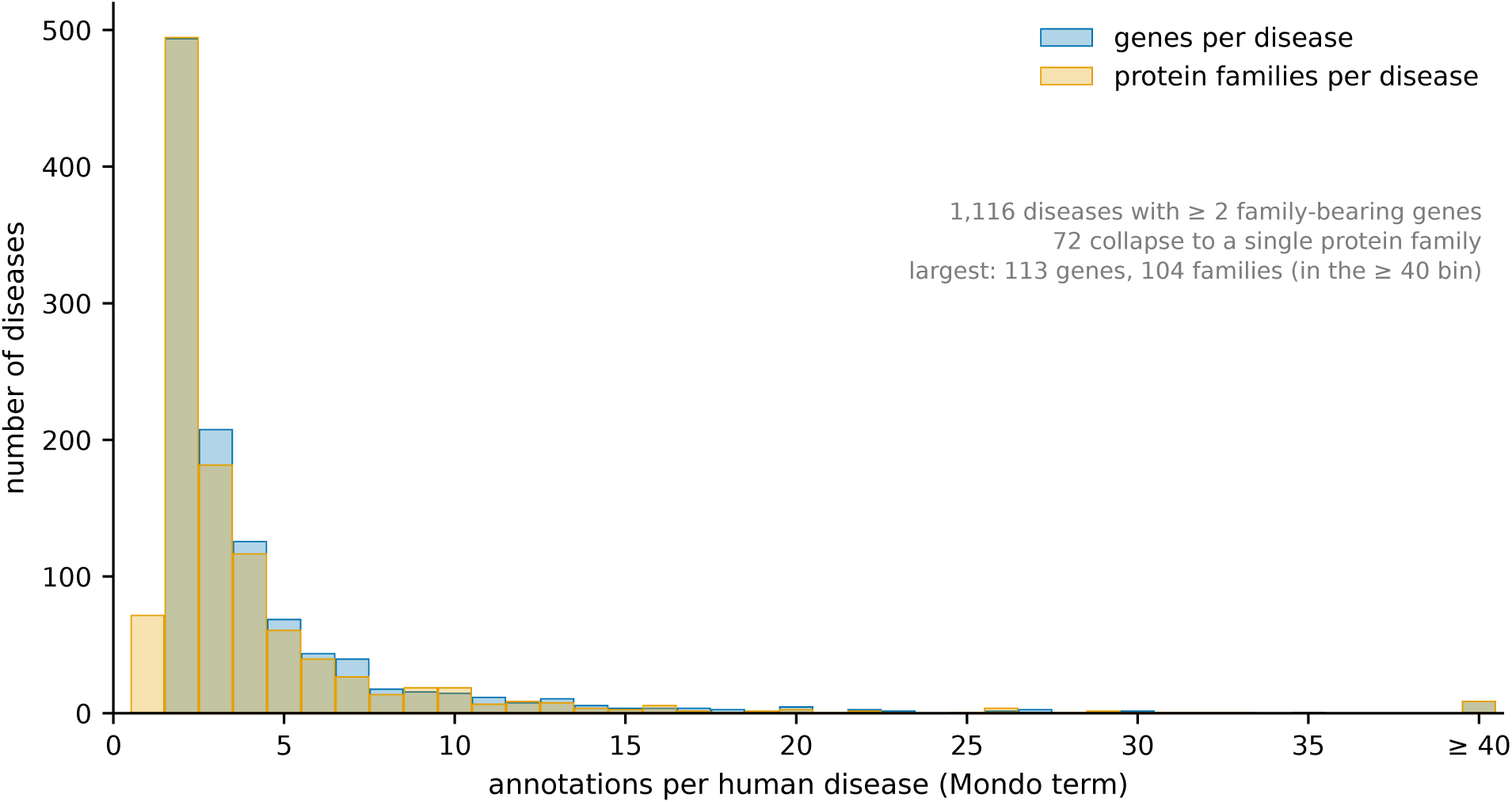
Distribution of the number of protein families or genes associated with each MONDO disease term. Gene-disease networks remain stable when collapsed to the orthogroup level, with only 72 disease nodes collapsing down to a single protein family annotation from an original protein family annotation count of ≥ 2.

### Supplementary Note 3. False discovery rate for initial phenolog calculations

The number of phenolog comparisons to be made between any two species corresponds to the number of phenotypes in species-a multiplied by the number of phenotypes in species-b, where each phenotype must be linked to at least one orthologous gene between the two species. The number of initial phenolog calculations between human diseases and model species phenotypes is found in **Table S4**.

**Table S4.** Human gene-disease comparison between each model species phenotypes. Number of possible comparisons between human mondo disease terms and model species phenotype terms.

| Human disease (Species A) | Model species (Species B) | Comparable human diseases | Comparable phenotypes | Common orthologs | Pairwise comparisons |
| --- | --- | --- | --- | --- | --- |
| *Homo sapiens* | *Mus musculus* | 7,919 | 10,998 | 7,394 | 87,093,162 |
| *Homo sapiens* | *Danio rerio* | 7,725 | 26,040 | 6,479 | 201,159,000 |
| *Homo sapiens* | *Caenorhabditis elegans* | 5,925 | 1,707 | 3,931 | 10,113,975 |
| *Homo sapiens* | *Schizosaccharomyces pombe* | 3,072 | 3,646 | 2,301 | 11,200,512 |
| *Homo sapiens* | *Rattus norvegicus* | 7,879 | 1,156 | 7,296 | 9,108,124 |
| *Homo sapiens* | *Xenopus tropicalis* | 7,522 | 124 | 6,173 | 932,728 |
| *Homo sapiens* | *Dictyostelium discoideum* | 3,719 | 235 | 3,021 | 873,965 |

Given the sufficiently large number of comparisons being made between each pair of species, the false discovery rate correction methodology previously described in (McGary et al. 2010) is used. This method compares the phenologs computed from 1,000 randomized species gene-association networks. Note, that within this network randomization process, for any given gene-association (disease or phenotype), the number of genes originally associated with it will remain the same each trial, and a random set of genes is pulled from the common orthologs found between the two species. For a gene to be available to draw from the random pool, it must also be connected to a phenotype. Each trial computes the set of pairwise comparisons between the two species networks and the hypergeometric p-values are recorded across all comparisons and all trials. To obtain an estimated false discovery rate of 5% we compare the p-value distribution from the real phenolog calculations, to the distribution of the p-values taken from the randomized trials. We find the p-value at which 5% or less of the real data overlaps with the randomized data. In the example given in **Fig. S3**, we show the log scaled histogram of p-values obtained by comparing all human diseases against all mouse phenotype terms (“real”), along with the 1,000 randomized versions of these calculations. Note, we first pool all 1,000 random trials together, create the histogram, then divide the y-values obtained (number of data-points within a given x-axis bin) by 1,000 to set the randomized trial data on the same scale as our real calculations. In this example we find 51,766 real “phenologs” with a p-value ≤ 2.7 × 10^−4^, in which we could expect 3.8% to be false positives given the overlap found with the randomized background distribution. The next higher p-value would push that overlap beyond the allowed 5% so we stop at 3.8%. This becomes our “significance” threshold for determining whether or not to include a disease-phenotype comparison in the set of data used downstream. The resulting number of disease-phenotype comparisons passing each species threshold can be found in **(Fig. 2C)**.

**Figure S3.**
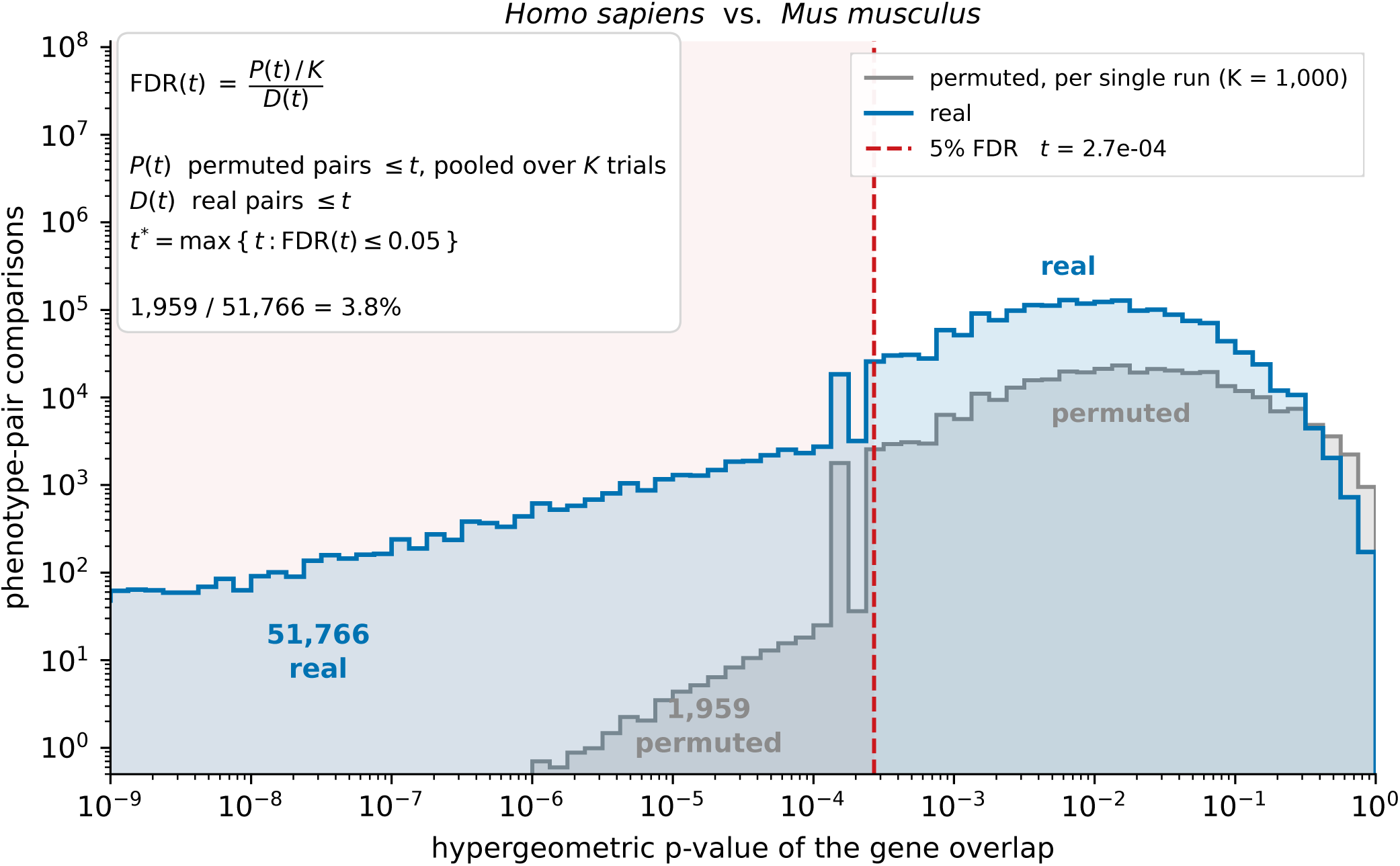
A visualization of how we compute the cut-off p-value for our false discovery rate corrections. In this figure we present the distribution of p-values for the disease-phenotype linkages from real data and p-values from one thousand permuted trials between human and mouse. The dashed red line shows the maximum point where our goal of having at least twenty times more real pairs than permuted pairs is achieved (i.e. 5% estimated FDR).

**Figure S4.**
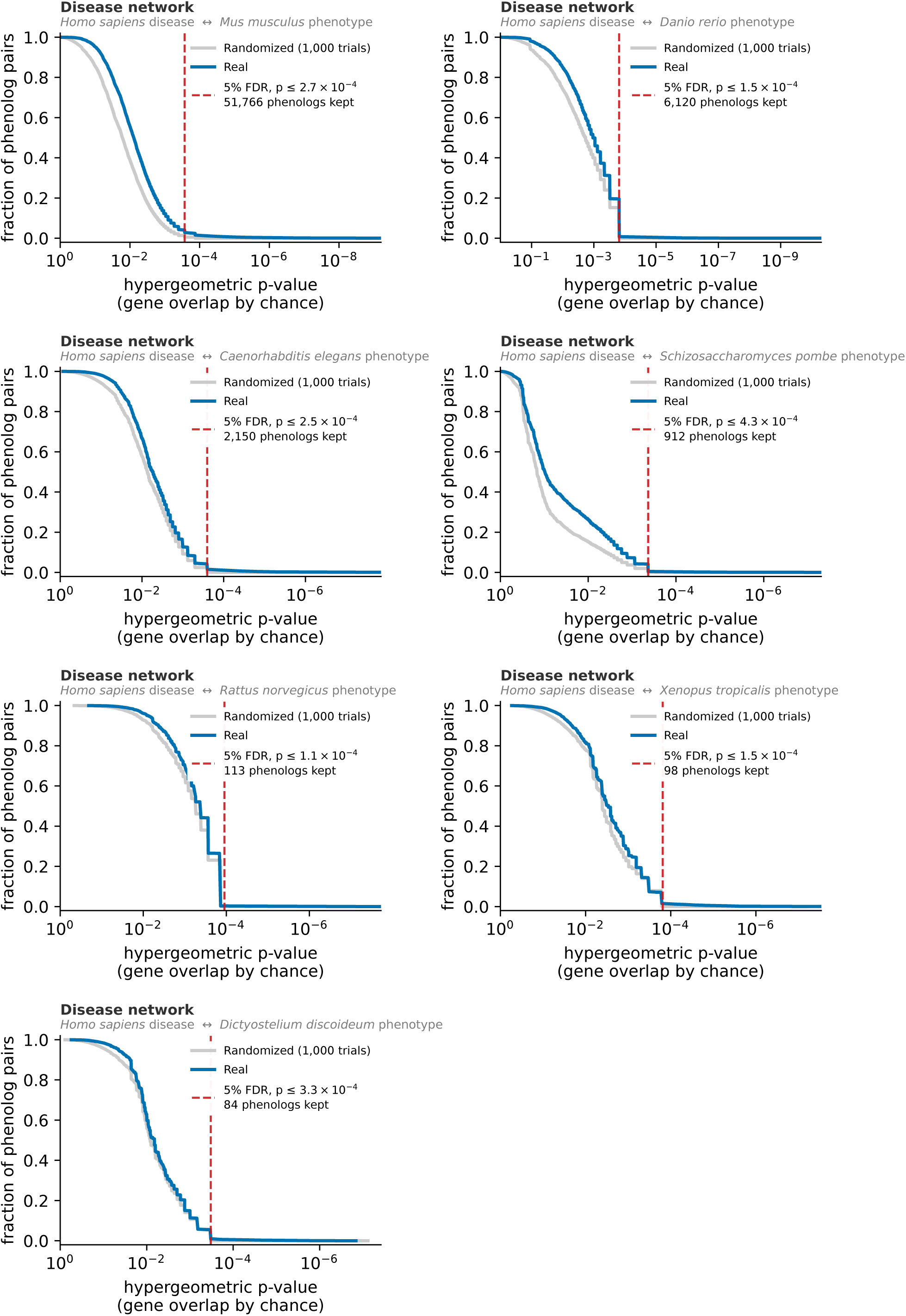
Phenolog significance determination, Human gene-disease network. Results of the 1,000 random trials between randomized human gene-disease and model species gene-phenotype networks for each species within our comparison set. Phenolog p-values are pooled across all 1,000 random trials within a given model species comparison, and then sorted (for comparisons where there is ≥ 1 gene/ortholog overlapping). We highlight with the red dashed line the p-values used in this study for each species representing an estimated 5% false discovery rate in accordance with work prior (McGary et al. 2010).

**Figure S5.**
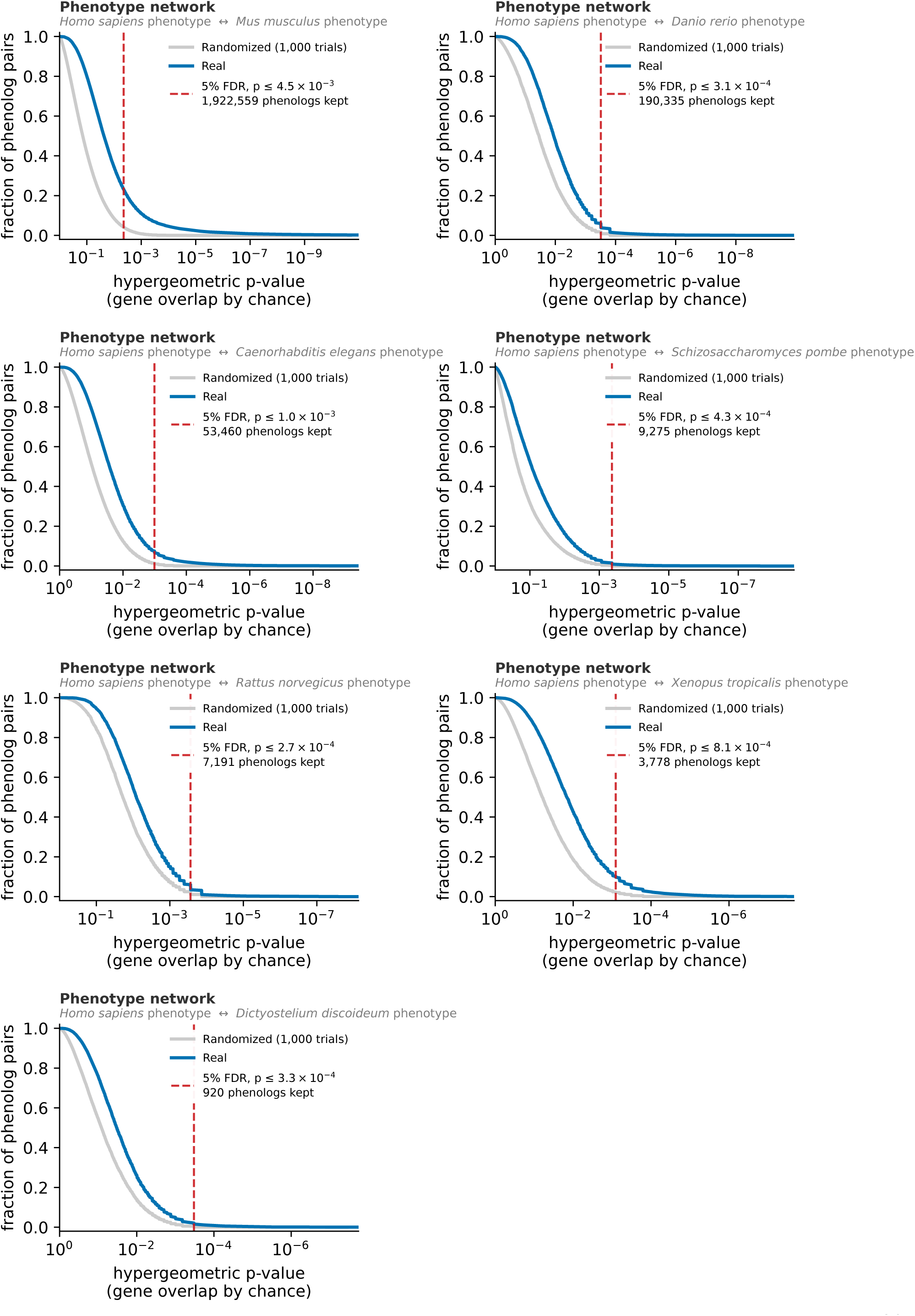
Phenolog significance determination, Human gene-phenotype network. Results of the 1,000 random trials between randomized human gene-phenotype and model species gene-phenotype networks for each species within our comparison set. Phenolog p-values are pooled across all 1,000 random trials within a given model species comparison, and then sorted (for comparisons where there is ≥ 1 gene/ortholog overlapping). We highlight with the red dashed line the p-values used in this study for each species representing an estimated 5% false discovery rate in accordance with work prior (McGary et al. 2010)

### Supplementary Note 4. Naive Bayes gene ranking method

To integrate information across multiple “phenologs”, for a single human disease, (Woods et al. 2013) used a matrix based formalization to help describe the problem where *X* is a set of gene-phenotype predictions, and *X_ij_* describes a value associating gene *i* to phenotype *j*. Here, we are using human diseases as the “phenotype” columns, and HGNC identifiers as the “gene” rows. In this way, one is able to rank how strongly any given *gene i* is associated with *disease j* across all genes and diseases. To compute *X_ij_* both (McGary et al. 2010) and (Woods et al. 2013) use the following equation, which they describe as the naïve Bayes scheme:

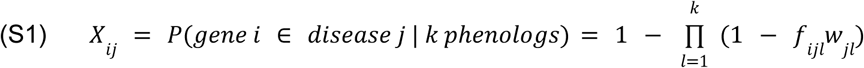

where

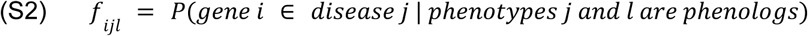

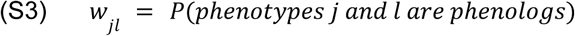

Here, *w_jl_* is 1 - hypergeometric p-value that was used to compute the initial association (McGary et al. 2010). For *f_ijl_*, we use *v*/*n*, which is the “fraction of the number of genes common to both phenotypes *j* and *l* over the number of genes known to be involved in phenotype *j”* (Woods et al. 2013). In our case, phenotype *j* is a human disease, and phenotype *l* is a cross species phenotype. In this way, we are computing the probability of the phenolog association being true multiplied by the approximate probability of a single gene-disease association already being true for a particular phenolog pairing. Subtracting from 1 then gives the probability of that gene-disease pairing being “false”. Multiplying these probabilities together across k phenologs and further subtracting from 1 gives the probability of that gene-disease pairing being true. We will also refer to this method the naive Bayes (nb) metric as previously described.

### Supplementary Note 5. Hypergeometric gene ranking method

**Figure S6.**
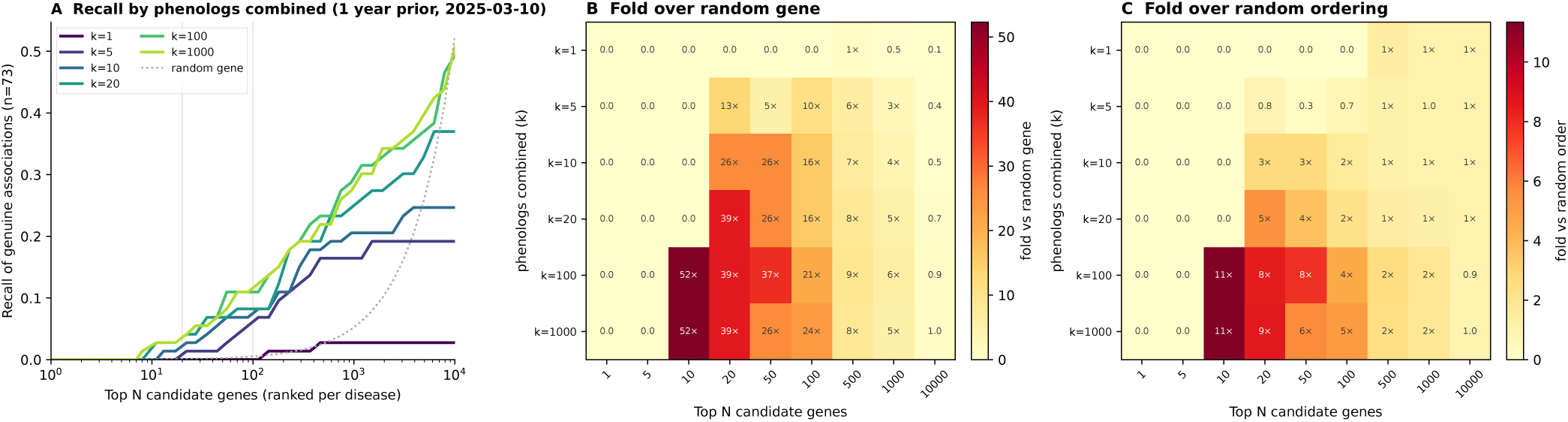
Parameter sweep of the one-year temporal validation across phenologs combined (k) and shortlist depth (N). All panels use hg rankings built from Monarch KG release 2025-03-10 and scored against the 73 genuine gene-disease pairs added by release 2026-03-06 (Table 2), as in Fig. 3. (A) Recall of the 73 pairs within each disease’s top N candidate genes for k = 1, 5, 10, 20, 100 and 1,000 phenologs combined per disease, one curve per k. The dotted line is the recall expected when N genes are drawn at random from the U human genes carrying an ortholog edge to a modeled species, min(N, U)/U. Faint vertical lines mark N = 20 and N = 100. (B) Fold enrichment over that random-gene expectation for every combination of k (rows) and N (columns); each cell is the recall at N divided by min(N, U)/U, printed in the cell (values of one or more carry a multiplication sign). (C) Fold enrichment over a random ordering of each disease’s own candidate list: the recall at N divided by the recall expected if the candidate genes surfaced for a disease were ranked at random, which for a pair whose disease has M candidates is min(N, M)/M. Panel B measures the gain from restricting a disease to its phenolog-derived candidates and ranking them; panel C isolates the gain from the ranking alone.

We introduce a new method for integrating information across top k-associated phenologs. For this new formulation of the gene-disease association matrix *X*, we define each gene-disease association value by computing the upper-tail probability of the hypergeometric distribution of the gene’s occurrence within a disease’s top k phenologs P(X≥c). This is implemented using the survival function in Python’s SciPy as,

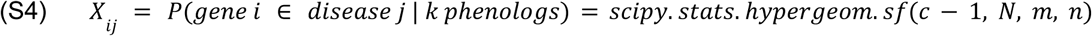

where *n* is the total number of genes found connected to any of the top k ranked phenotypes (phenologs). Or more formally:

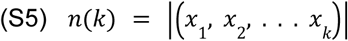

where *x* is a gene belonging to a phenotype within the top k ranked phenologs. In our metric formulation, *n* can be thought of as the number of times we draw a gene prediction given our initial pool of candidate phenotypes found to be associated with a disease. The number of times each gene occurs within our background network is *m*. More specifically, *m* is the number of phenotypes the gene of interest is connected to across all species within our graph. *N* is the background total number of existing phenotype-gene edges where each gene must have an ortholog in human (or the base species one is using for comparison). The number of times each gene shows up within our top ranked k phenologs is used as *c*. We can then ask what is the probability of gene *i* showing up >= *c* times given its background count within the network *m*, *N* total associations we could potentially draw from, and *n* genes we originally pulled in from our k top ranked phenologs.

### Supplementary Note 6. Temporal evaluation parameter sweep

The analysis leading to (**Table 2**) identified 73 genuine new gene-disease pairs added to the Monarch KG between releases 2025-03-10 and 2026-03-06, spanning 43 diseases. As in (**Fig. 3A**), we rank each of these diseases’ candidate genes from the earlier release and ask where the new genes fall, here across a grid of k, the number of top phenologs combined per disease, and N, the candidate gene list depth (**Fig. S6**). Enrichment over a randomly drawn gene is highest when only considering the top candidates, where recall is still small, so the practical window appears to be roughly the top 20 to 100 candidates, which retains double-digit fold enrichment together with meaningful recall, and enrichment still eightfold or more out to the top 500 (**Fig. S6A,B**). The fold enrichment over a random ordering of each disease’s own candidate list is smaller, 11-fold at N = 10 and 4 to 5-fold at N = 100, and is the stricter measure of what the ranking itself adds (**Fig. S6C**). The permutation p-value reported in the main text tests this stricter null, comparing the observed recall at N with 5,000 random orderings of each disease’s candidate gene list, so the smallest attainable value is 2 × 10⁻⁴. Within this window hg and nb are statistically indistinguishable (p = 0.32; **Fig. 3A**), as expected given the fact they score from the same pool of surfaced genes. This is also consistent with the parity between nb and hg when ranking candidates within a disease under leave-one-out evaluation (**Supplementary Note 7**).

### Supplementary Note 7. Leave-one-out performance evaluation

We adapt the orthogroup-based leave-one-out cross validation of (Woods et al. 2013) to ask whether known gene-disease associations within the Monarch KG can be recovered once they are hidden from the method, and how highly they rank when they are. For each of the 2,001 PANTHER protein families carrying at least one human disease annotation, we remove the family’s genes from the human gene-disease network, recompute the full set of phenologs, and rebuild the gene rankings with both the naive Bayes (nb) and hypergeometric (hg) schemes (Supplementary Notes 4 and 5) at k = 1, 5, 20 and 100 phenologs combined per disease. The whole family is withheld rather than a single gene because a paralog left behind in the same family would otherwise stand in for the hidden gene and make the test trivial. Ranks are then read at the gene level, so a family counts as recovered only through the specific genes that were annotated to the disease. Tied scores receive the average rank, and when a disease has more than one hidden gene we record the median of their ranks.

The resulting ranks are read in two directions (**Fig. S7A,B**). In the first, we take a disease and ask where its hidden genes fall among all candidate genes ranked for that disease (**Fig. S7A**). This is the candidate gene prediction task used throughout this work, and the comparison made by (Woods et al. 2013). In the second, we take a hidden gene and ask where its true disease falls among all diseases the gene is ranked for (**Fig. S7B**). This transposed view is only meaningful if scores can be compared across diseases, which the nb scheme was never designed for. A disease enters a cell of either heatmap when its median rank is at or below the column threshold, and the total column counts the diseases whose hidden genes were surfaced at any rank. A disease must retain at least one other annotated family for any phenolog to be found, so diseases annotated to a single protein family (**Fig. S2**) cannot be evaluated, and coverage grows with k as more phenologs contribute candidates (168, 274, 322 and 332 diseases at k = 1, 5, 20 and 100).

Within a disease the two schemes are close to interchangeable (Fig. S7A). Both draw on the same candidate pool, so the total column is identical, and the counts differ by at most 24 diseases at any threshold, with the two schemes trading places from one cut-off to the next and nb marginally ahead at the looser ones. Across diseases the picture changes completely (**Fig. S7B**). At k = 20, hg places the true disease first for 18 diseases and within the top 20 for 165, whereas nb manages 0 and 12. nb also loses resolution as more phenologs are combined, consistent with its scores saturating towards one as the product in (**Eq. S1**) accumulates terms: the diseases it places within the top 20 fall from 119 at k = 1 to 4 at k = 100, while hg holds between 147 and 165 at every k. This is the property that allows hg candidates to be pooled into a single ranked list across diseases (**Fig. 3B**).

Panels C and D complement the leave-one-out analysis with a direct comparison of how the two schemes order the same candidates, computed from the complete model at k = 100 across the 414 diseases with at least four known genes. No genes are withheld here, so a disease’s own annotations help find its phenologs and the absolute recall is optimistic, but both schemes are measured on identical candidate lists. hg recovers a larger fraction of each disease’s known genes at every shortlist size (45% versus 39% within the top 20; **Fig. S7C**), and its per-disease area under the precision-recall curve is higher in 278 of 414 diseases (67%), by a median of 0.026 (**Fig. S7D**). Taken together, hg matches nb at ranking candidates within a disease, improves precision at the top of each list, and is the only one of the two that can be used to rank candidates across diseases.

**Figure S7.**
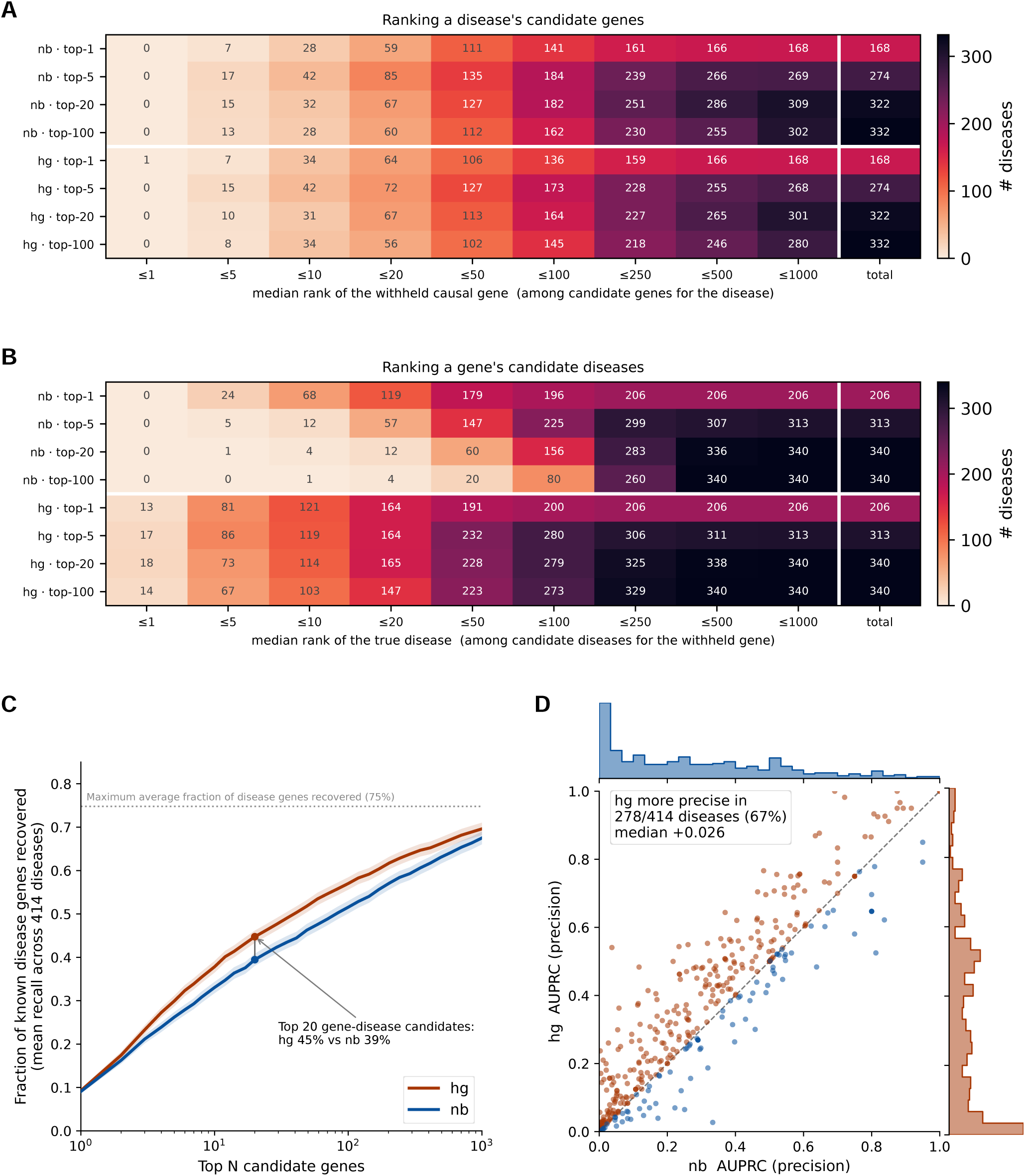
Leave-one-out recovery of known gene-disease associations and the precision of the two gene ranking schemes. (A) Ranking a disease’s candidate genes. Each row is one ranking scheme (nb or hg) at one setting of k, the number of top phenologs combined per disease (1, 5, 20, 100). For each of the 2,001 PANTHER protein families carrying a human disease annotation, the family’s genes are removed from the human gene-disease network, phenologs and gene rankings are recomputed, and the rank of each withheld gene is read from the ranked candidate list of every disease it was annotated to; tied scores receive the average rank. A disease’s value is the median rank over its withheld genes. Cells count the diseases whose median rank is at or below the column threshold, and the total column counts the diseases whose withheld genes appear anywhere in the candidate list. The colour scale runs from 0 to the largest total (332 in A, 340 in B). (B) Ranking a gene’s candidate diseases. The same leave-one-out runs read in the other direction: the diseases ranked for each withheld gene are sorted by score and the rank of the gene’s true disease is recorded, then summarized per disease and counted as in A. This direction requires scores that are comparable across diseases. (C) Recall of known disease genes by the complete model (no genes withheld; k = 100) across the 414 diseases with at least four known genes carrying an ortholog in a modeled species. Curves give the mean fraction of a disease’s known genes found within its top N candidates (shading, ± 1 s.e.m.); the dotted line is the mean fraction of known genes present anywhere in the candidate lists (75%), identical for both schemes because they rank the same candidates. At N = 20, hg recovers 45% and nb 39%. (D) Per-disease area under the precision-recall curve (AUPRC) for the same 414 diseases, nb on the x axis and hg on the y axis, with marginal histograms. Orange points lie on or above the diagonal (hg at least as precise) and blue points below it. hg is more precise in 278 of 414 diseases (67%), by a median of 0.026.

## Notes

https://monarch-initiative.github.io/monarch-phenologs/

### Competing Interest Statement

The authors have declared no competing interest.

